# Drifting codes within a stable coding scheme for working memory

**DOI:** 10.1101/714311

**Authors:** M. J. Wolff, J. Jochim, E. G. Akyürek, T. J. Buschman, M. G. Stokes

## Abstract

Working memory (WM) is important to maintain information over short time periods to provide some stability in a constantly changing environment. However, brain activity is inherently dynamic, raising a challenge for maintaining stable mental states. To investigate the relationship between WM stability and neural dynamics, we used electroencephalography to measure the neural response to impulse stimuli during a WM delay. Multivariate pattern analysis revealed representations were both stable and dynamic: there was a clear difference in neural states between time-specific impulse responses, reflecting dynamic changes, yet the coding scheme for memorized orientations was stable. This suggests that a stable subcomponent in WM enables stable maintenance within a dynamic system. A stable coding scheme simplifies readout for WM-guided behaviour, whereas the low-dimensional dynamic component could provide additional temporal information. Despite having a stable subspace, WM is clearly not perfect – memory performance still degrades over time. Indeed, we find that even within the stable coding scheme, memories drift during maintenance. When averaged across trials, such drift contributes to the width of the error distribution.

## Introduction

Neural activity is highly dynamic, yet often we need to hold information in mind in a stable state to guide ongoing behaviour. Working memory is a core cognitive function that provides a stable platform for guiding behaviour according to time extended goals; however, it remains unclear how such stable cognitive states emerge from a dynamic neural system.

At one extreme, WM could effectively pause the inherent dynamics by falling into a stable attractor (e.g., 1,2). This solution has been well-studied, and provides a simple readout of memory content irrespective of time (i.e., memory delay). However, more dynamic models have also been suggested. For example, in a recent hybrid model, stable attractor dynamic coexist with a low-dimensional, time varying component (3,4); see Fig. 1A for model schematics). This permits some dynamic activity, whilst also maintaining a fixed coding relationship of WM content over time (5). As in the original stable attractor model, the coding scheme is stable over time, permitting easy and unambiguous WM read out by downstream systems, regardless of maintenance duration (6). Finally, it is also possible to maintain stable information in a richer dynamical system (e.g., 7). Although the relationship between activity pattern and memory content changes over time, the representational geometry could remain relatively constant (5). Such dynamics emerge naturally in a recurrent network, and provide rich information about the previous input, and elapsed time (8), but necessarily entail a more complex readout strategy (time-specific decoders or a high-dimensional classifier that finds a high-dimensional hyperplane that separates memory condition for all time points - (9)).

**Figure 1.**
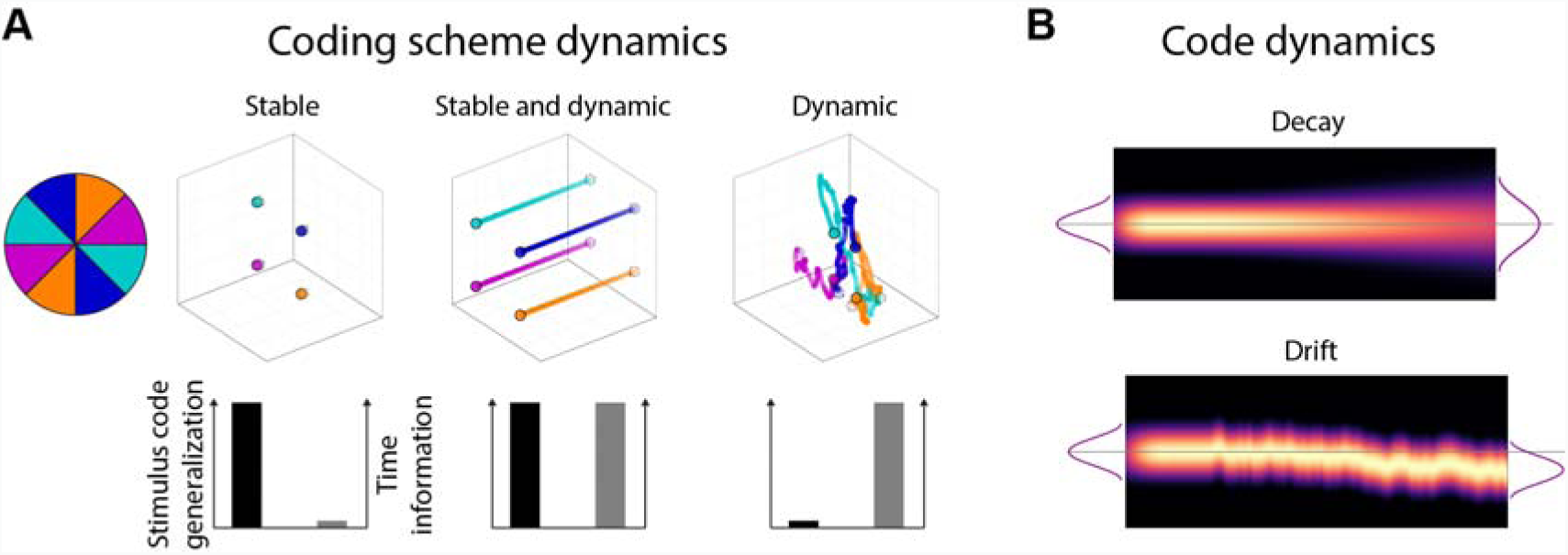
Model predictions. (A) The relationship between the neural coding scheme of orientations (colours) in WM over time. Left: A stable coding scheme within a stable neural population. Middle: A stable coding scheme within a dynamic neural population. Right: A dynamically changing coding scheme. (B) The fidelity of the population code in WM over time. Top: The code decays and becomes less specific over time, leading to random errors during read-out. Bottom: The code drifts along the feature dimension, leading to a still sharp, but shifted code during read-out.

Although all models seek to account for stable WM representation, it is also important to note that maintenance in WM is far from perfect. In particular, WM performance decreases over time. (10), which could be ascribed to two different mechanisms (Fig. 1B). On the one hand, the neural representation could simply degrade over time, either due to an overall decrease in WM specific neural activity, or through a general broadening of the neural representation (11). In this framework, the distribution of recall error reflects sampling from a broad underlying distribution. On the other hand, the neural representation of WM content might gradually drift along the feature dimension as a result of the accumulating effect of random shifts due to noise (12). Even if the underlying neural representation remains sharp, variance in the mean over trials results in a relative broad distribution of errors over trials.

Computational modelling based on behavioural recall errors from WM tasks with varying set-sizes and maintenance periods predict a drift for colours and orientations maintained in WM (13,14). At the neural level, evidence for drift has been found in the neural population code in monkey prefrontal cortex during a spatial WM task (15), where trial-wise shifts in the neural tuning profile predicted if recall error was clockwise or counter-clockwise relative to the correct location. Recently, a human fMRI study has found that delay activity reflected the probe stimulus more when participants erroneously concluded that it matched the memory item (16), which is consistent with the drift account.

Tracking these neural dynamics of non-spatial neural representations, which are not related to spatial attention or motor planning, is not trivial in humans. Previously we found that the presentation of a simple impulse stimulus (task-relevant visual input) presented during the maintenance period of visual information in WM results in a neural response that reflects non-spatial WM content (17,18). Here we extend this approach to track WM dynamics. In the current study we developed a paradigm to test the stability (and/or dynamics) of WM neural states and the consequence for readout by “pinging” the neural representation of orientations at specific time-points during maintenance.

We found that the coding scheme remained stable during the maintenance period, even-though maintenance time was coded in an additional low-dimensional axis. We furthermore found that the neural representation of orientations drifts in WM. This was reflected in a shift of the reconstructed orientation towards the end of the maintenance period that predicted behaviour.

## Methods

### Participants

Twenty-six healthy adults (17 female, mean age 25.8 years, range 20-42 years) were included in all analyses. Four additional participants were excluded during preprocessing due to excessive eye-movements (more than 30% of trials contaminated). Participants received monetary compensation (£10 an hour) for participation and gave written informed consent. The experiment was approved by the Central University Research Ethics Committee of the University of Oxford.

### Apparatus and stimuli

The experimental stimuli were generated and controlled by Psychtoolbox (19), a freely available Matlab extension. Visual stimuli were presented on a 23-inch (58.42 cm) screen running at 100 Hz and a resolution of 1,920 by 1,080. Viewing distance was set at 64 cm. A Microsoft Xbox 360 controller was used for response input by the participants.

A grey background (RGB = 128, 128, 128; 20.5 cd/m^2^) was maintained throughout the experiment. A black fixation dot with a white outline (0.242°) was presented in the centre of the screen throughout all trials. Memory items and the probe were sine-wave gratings presented at 20% contrast, with a diameter of 8.51° and spatial frequency of 0.65 cycles per degree, with randomised phase within and across trials. Memory items were presented at 6.08° eccentricity. The rotation of memory items and probe were randomized individually for each trial. The impulse stimulus was a single white circle, with a diameter of 20.67°, presented at the centre of the screen. The retro-cue was two arrowheads pointing right (>>) or left (<<), and was 1.58° wide. A coloured circle (3.4°) was used for feedback. Its colour depended dynamically on the precision of recall, ranging from red (more than 90 degrees error) to green (0 degrees error). A pure tone also provided feedback on recall accuracy after each response, ranging from 200 Hz (more than 90 degrees error) to 1,100 Hz (0 degrees error).

### Procedure

Participants participated in a free-recall, retro-cue visual WM task. Each trial began with the fixation dot. After 1,000 ms the memory array was presented for 200 ms. After a 400 ms delay, the retro-cue was presented for 100 ms, indicating which of the previously two items would be tested, rendering the other item irrelevant. The first impulse stimulus was presented for 100 ms, 900 ms after the offset of the retro-cue. After a delay of 700 ms, the second impulse stimulus was presented for 100 ms. After another delay of 700 ms the probe was presented. Participants used the left joystick on the controller with the left thumb to rotate the orientation of the probe until it best reflected the memorized orientation, and confirmed their answer by pressing the “x” button on the controller with the right thumb. Note that one complete rotation of the joystick corresponded to 0.58 of a rotation of the probe. In conjunction with the fact that the probe was randomly orientated on each trial, it was impossible for participants to plan the rotation beforehand or memorize the direction of the joystick instead of the orientation of the memory item. Accuracy feedback was given immediately after the response where both the coloured circle and tone were presented simultaneously. Each participant completed 1,100 trials in total, over a course of approximately 135 minutes, including breaks. See Figure 2A for a trial schematic.

**Figure 2.**
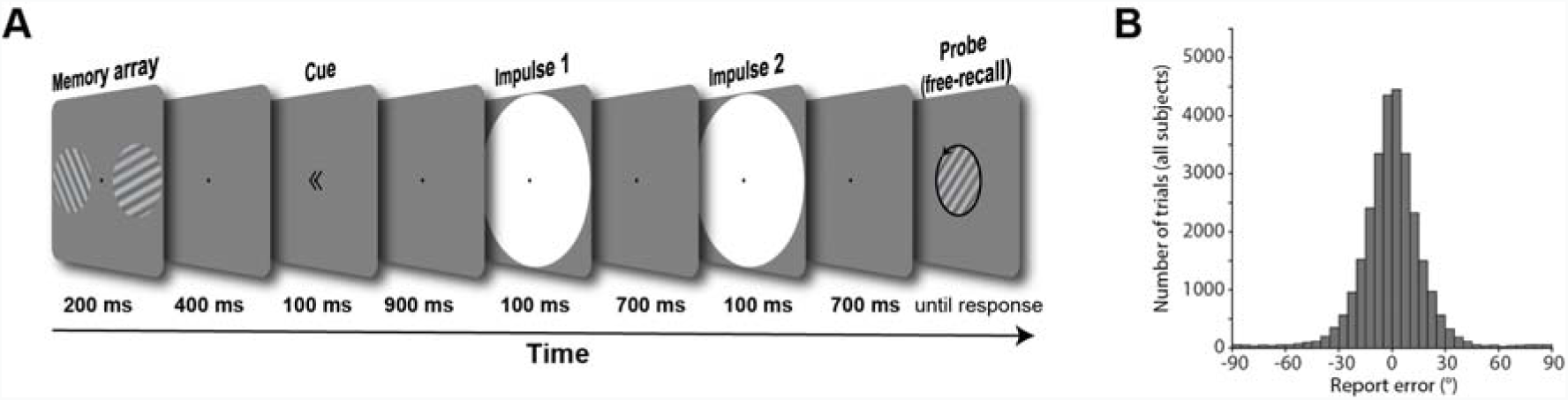
Trial schematic and behavioural results **(A)** Two randomly orientated grating stimuli were presented laterally. A retro-cue then indicated which of those two would be tested at the end of the trial. Two impulses (white circles) were serially presented in the subsequent delay period. At the end of the trial a randomly oriented probe grating was presented in the centre of the screen, and participants were instructed to rotate this probe until it reflected the cued orientation. **(B)** Report errors of all trials across all subjects.

### EEG acquisition

EEG was acquired with 61 Ag/AgCl sintered electrodes (EasyCap, Herrsching, Germany) laid out according to the extended international 10–20 system and recorded at 1,000 Hz using Curry 7 software (Compumedics NeuroScan, Charlotte, NC). The anterior midline frontal electrodes (AFz) was used as the ground. Bipolar electrooculography (EOG) was recorded from electrodes placed above and below the right eye and the temples. The impedances were kept below 5 kΩ. The EEG was referenced to the right mastoid during acquisition.

### EEG preprocessing

Offline, the EEG signal was re-referenced to the average of both mastoids, down-sampled to 500 Hz, and bandpass filtered (0.1 Hz high-pass and 40 Hz low-pass) using EEGLAB (20). The continuous data was epoched relative to the memory array onset (−500 ms to 3,600 ms) before independent component analysis (21) was applied. Components related to eye-blinks were subsequently removed. The data was then epoched relative to memory array onset and the two impulse onsets (0 ms to 400 ms), and trials were individually inspected. Trials with saccadic eye movements, visually identified from the electrooculography, and trials with non-archetypical artefacts, visually identified from the EEG, in the memory array epoch and in either impulse epoch were removed from all subsequent analyses. Furthermore, trials where the report error was 3 circular standard deviations from the participant’s mean response error were also excluded from EEG analyses to remove trials that likely represent complete guesses (22). This lead to the removal of *M* = 2.3% (*SD* = 1.2%) trials due to inaccurate report trials, in addition to the *M* = 3.52 % (*SD* = 4.21%) and *M* = 5% (*SD* = 5.2%) of trials removed due to eye-movements and non-archetypical EEG artefacts from the memory array and impulse epochs, respectively.

While MVPA on electrophysiological data is usually performed on each time-point separately, taking advantage of the highly dynamic waveform of evoked responses in EEG by pooling information multivariately over electrodes as well as time can improve decoding accuracy, at the expense of temporal resolution (23,24). Since the previously reported WM-dependent impulse response reflects the interaction of the WM state at the time of stimulation and does not reflect continuous delay activity, we treat the impulse responses as discrete events in the current study. Thus, the whole time window of interest relative to impulse onsets (100 to 400 ms) from the 17 posterior channels was included in the analysis. The time window was based on previous, time-resolved findings, which showed that the WM-dependent neural response from a 100 ms impulse (as used in the current study) is largely confined to this window (18). In the current study, instead of decoding at each time-point separately, information was pooled across the whole time-window. The mean activity level within each time window of each channel was first removed, thus normalizing the voltage fluctuations and isolating the dynamic, impulse-evoked neural signal from more stable brain states. The time-window was then down-sampled by taking the average every 10 ms, thus resulting in 50 values per channel, each of which was treated as a separate dimension in the subsequent multivariate analysis (850 in total). This data format was used on all subsequent MVPA analyses, unless explicitly mentioned otherwise. The same approach over the same time window of interest was used in our previous study (25).

### Orientation reconstruction

We computed the mahalanobis distances as a function of orientation difference to reconstruct grating orientations (18). The following procedure was performed separately for items that were presented on the left and right side. Since the grating orientations were determined randomly on a trial-by-trial basis and the resulting orientation distribution across trials was unbalanced, we used a k-fold procedure with subsampling to ensure unbiased decoding. Trials were first assigned the closest of 16 orientations (variable, see below) which were then randomly split into 8 folds using stratified sampling. Using cross-validation, the train trials in 7 folds were used to compute the covariance matrix using a shrinkage estimator (26). The number of trials of each orientation bin were equalized by randomly subsampling the minimum number of trials in any bin. The subsampled trials of each angle bin were then averaged. To pool information across similar orientations, the average bins were convolved with a half cosine basis set raised to the 15^th^ power (27–29). The mahalanobis distances between each trial of the left-out test fold and the averaged and basis-weighted angle bins were computed and mean-centred across the 16 distances to normalize. This was repeated for all test and train fold combinations. To get reliable estimates, the above procedure was repeated 100 times (random folds and subsamples each time), separately for eight orientation spaces (0° to 168.75°, 1.40625° to 170.1563°, 2.8125° to 171.5625°, 4.2188° to 172.9688°, 5.625° to 174.375°, 7.0313° to 175.7813°, 8.4375° to 177.1875°, 9.8438° to 178.5938°, each in steps of 11.25°). For each trial we thus obtained 800 samples for each of the 16 mahalanobis distances. The distances were averaged across the samples of each trial and ordered as a function of orientation difference. The resulting “tuning curve” was summarized into a single value (i.e., “decoding accuracy”) by computing the cosine vector mean of the tuning curve (18), where a positive value suggests a higher pattern similarity between similar orientations than between dissimilar orientations. The approach was the same for the reanalysis of (17).

We also repeated the above analysis iteratively for a subset of electrodes in a searchlight analysis across all 61 electrodes. In each iteration, the “current” as well as the closest two neighbouring electrodes were included in the analysis (similar as in 30) The freely available MATLAB extension fieldtrip (31) was used to visualise the decoding topographies. Note that the topographies were flipped, such that the left represents the ipsilateral and the right the contralateral side relative to stimulus presentation side.

### Orientation code generalization

To test cross-generalization between impulses, instead of training and testing within the same time-window, the train folds were taken from impulse 1, and the test fold from impulse 2, and vice versa. The analysis was otherwise exactly as described above.

To test cross-generalization between presented locations, the classifier was similarly trained on trials where the item was presented on the left, and tested on the right, and vice versa. Since left and right trials were independent trial sets, cross-validation does not apply. However, to ensure a balanced training set, the number of trials of each orientation bin were nevertheless equalized by subsampling (as described above), and this approach was repeated 100 times.

The cross-generalization of the orientation code between impulse onsets in (17) was tested with the same analyses as the location cross-generalization described in the paragraph above: The classifier was trained on the early onset condition, and tested on the late-onset condition, and vice versa, while making sure that the training set is balanced through random subsampling.

### Impulse/time and location decoding

To decode the difference of the evoked neural responses between impulses, we used a leave-one-out approach. The mahalanobis distances between the signals from a single trial from both impulse epochs and the average signal of all other trials of each impulse epoch were computed. The covariance matrix was computed by concatenating the trials of each impulse (excluding the left-out trial). The average difference of same impulse distances were subsequently subtracted from different impulse distances, such that a positive distance difference indicates more similarity between same than different impulses. To convert the distance difference into trial wise decoding accuracy, positive distance difference were simply converted into “hits” (1) and negative into “misses” (0). The percentage of correctly classified impulses were subsequently compared to chance performance (50%).

The presentation side and impulse onset (in (17)) was decoded using 8-fold cross-validation, where the distance difference between different and same location/onset was computed for each trial, which were then converted to “hits” and “misses”.

### Visualization of the spatial, temporal, and orientation code

To explore and visualize the relationship between the location or impulse/time code and the orientation code in state space (see Fig. 1A for different predictions), we used classical multidimensional scaling (MDS) of the mahalanobis distances between the average signal of trials belonging to one of four orientation bins (0° to 45°, 45° to 90°, 90° to 135°, 135° to 180°) and location (left/right) or time (impulse 1/impulse2).

For the visualization of the code across impulse/time, distances were computed separately for left and right trials, before taking the average. Within each orientation bin, the data of half of the trials were taken from impulse 1, and the data of the other half from impulse 2 (determined randomly). The number of trials within each orientation of each impulse were equalized through random subsampling before averaging. The mahalanobis distances between both orientation and impulses were then computed using the covariance matrix estimated from all trials of both impulses. This was repeated 100 times (for each side), randomly subsampling and splitting trials between impulses each time and then taking the average across all iterations.

For the visualization of the code across space, the data of each trial were first averaged across impulses. The number of trials of orientation bins (same as above) of each location were equalized through random subsampling. The mahalanobis distances of the average of each bin within each location condition were computed using covariance estimated from all left and right trials. This was repeated 100 times, before taking the average across all iterations.

For the code across impulse onset/time visualization of the data from (17), the same procedure as in the paragraph above was used, but instead of visualizing the stimulus code between locations, it was visualized between impulse onsets (−30 ms, +30 ms).

### Relationship between behaviour and the neural representation of the WM item

We were interested if imprecise reports that are clockwise (CW) or counter-clockwise (CCW) relative to the actual orientation are accompanied by a corresponding shift of the neural representation in WM (see Fig. 1B for model schematics). We used two approaches to test for such a shift (Fig. 5A & 6A).

First, the trial-wise pattern similarities as a function of orientation differences (as obtained from the orientation-reconstruction approach described above) were averaged separately for all CW and CCW responses (Fig. 5A). Note that CW and CCW responses were defined relative to the median response error within each orientation bin. This ensures a balanced proportion of all orientations in CW and CCW trials, which is necessary to obtain meaningful orientation reconstructions. It furthermore removes the report bias away from cardinal angles in the current experiment (Suppl. fig. 1), similar to previous reports of orientation response biases (32), and thus isolates random from systematic report errors.

We used another approach that exaggerates the potential difference between CW and CCW trials and thus might be more sensitive to detect a shift. The data was first divided into CW and CCW trials using the same within orientation bin approach as described above. The classifier was then trained on CW trials, and tested on CCW trials, and vice versa (Fig. 6A). The orientation bins in the training set were balanced through random subsampling, and the procedure was repeated 100 times. Given an actual shift in the neural representation, the shift magnitude of the resulting orientation reconstruction of this method should be doubled, since both the testing data and the training data (the reference point) are shifted, but in opposite directions.

To improve orientation reconstruction from the impulse epochs, the classifier was trained on the averaged trials of both impulses, but tested separately on each impulse epoch individually. While training on both impulses improved orientation reconstruction, in particular for the second approach where only half of the trials are used for training, the shifts in orientation representations as a function of CW/CCW reports are qualitatively the same when training and testing within each impulse epoch separately (Fig. 5, 6, & Suppl. fig. 3).

### Statistical significance testing

To test for statistical significance of average decoding at the group level, the sign of the data of each participant was randomly flipped with a probability of 50% 100.000 times, and the resulting null-distribution was used to calculate the *p* value of the null hypothesis (no difference, chance decoding). Note that tests of within condition decoding (within presentation location, impulse/onset) were one-sided, since only positive decoding is plausible in those cases, whereas tests of cross-generalization between conditions were two-sided, since negative decoding is theoretically plausible in those cases. Comparisons of decodability between conditions/items were also two-sided.

The possible shift in representation towards the response was quantified and tested for statistical significance at the group level. The circular mean of the shifted average tuning curve (summarized such that a positive shift reflects a shift towards the response) was tested against 0. The tuning curve of each subject was flipped left to right with 0.5 probability, such that a subject’s positively shifted tuning curve would then be negatively shifted, before computing the circular mean of the resulting tuning curve averaged over all subjects 100.000 times. The resulting null distribution was used to obtain the p-value by calculating the proportion of permuted tuning curves with circular means more positive than the actual group-level circular mean. The test obtained p-value was one-sided, since we expected the shift of the neural representation of the orientation to be towards the response.

### Code and data availability

All data and custom Matlab scripts used to generate the results and figures of this manuscript will be made available upon peer-reviewed publication.

## Results

### Item and WM content-specific evoked responses during encoding and maintenance

The neural response elicited by the memory array contained parametric information about the presented orientations (*p* < 0.001, one-sided; Fig. 3, left).

**Figure 3.**
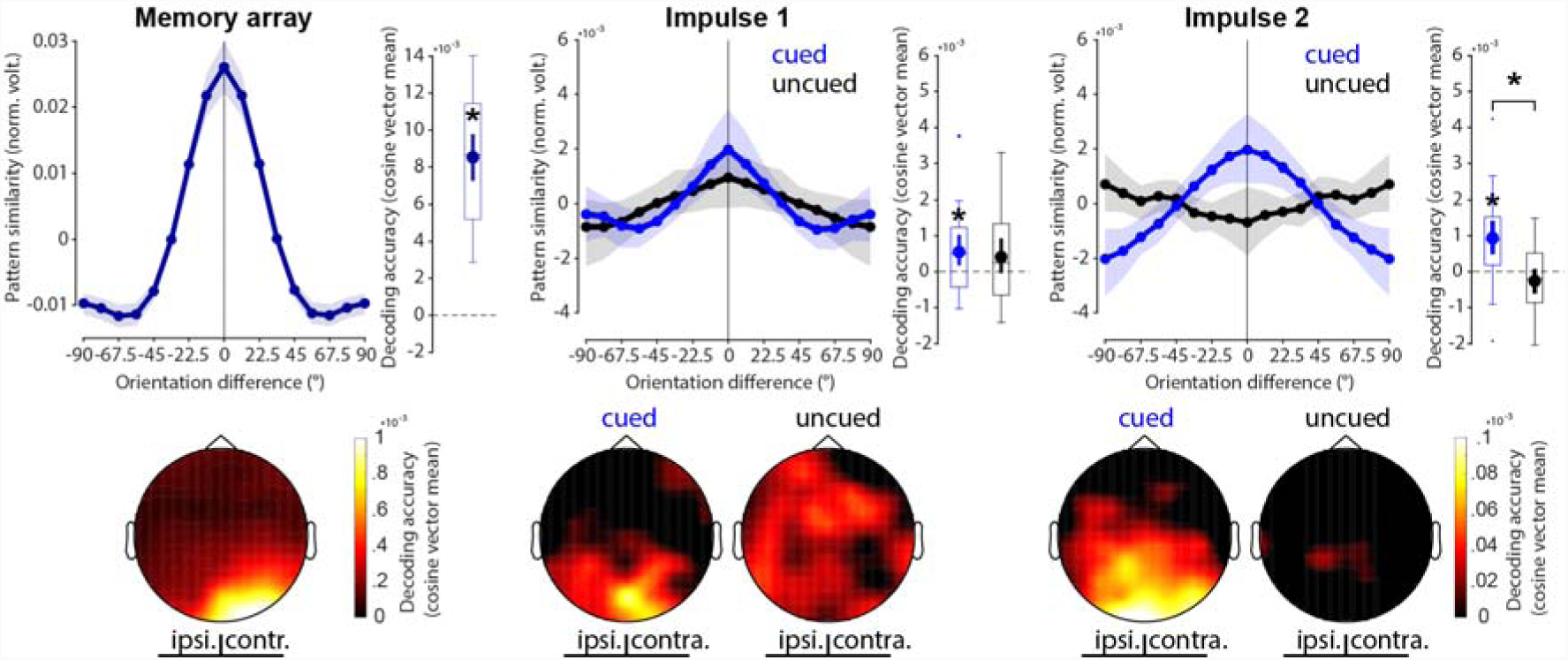
Decoding results. Top row: Normalized average pattern similarity (mean-centred, sign-reversed mahalanobis distance) of the evoked neural responses (100 to 400 ms relative to stimulus onset) as a function of orientation similarity, and decoding accuracy (cosine vector means of pattern similarities). Error shadings and error bars are 95 % C.I. of the mean. Centre lines of boxplots indicate the median; box outlines show 25th and 75th percentiles, and whiskers indicate 1.5x the interquartile range. Extreme values are shown separately (dots). Asterisks indicate significant decoding accuracies (p < 0.05, one-sided) or differences (p < 0.05, two-sided). Bottom row: Decoding topographies of the searchlight analysis.

The first impulse response contained statistically significant information about the cued item (*p* = 0.008, one sided), but not the uncued item, which failed to reach the statistical significance threshold (*p* = 0.057, one-sided). The difference between cued and uncued item decoding was not significant (*p* = 0.694, two-sided; Fig. 3, middle).

The decodability of the cued item was also significant at the second impulse response (*p* < 0.001, one-sided), while it was not of the uncued item (*p* = 0.919, one-sided). Notably, the decodability of the cued item was significantly higher than that of the uncued item (*p* = 0.002, two-sided; Fig. 3, right).

Overall, these results reflect previous findings (18) in that the impulse response reflects relevant information in WM, and that no longer relevant information leave no detectable trace in the WM network.

The decoding topographies highlight that most of the decodable signal came from posterior electrodes during both encoding and maintenance, and is therefore likely generated by the visual cortex. Notably, while contralateral electrodes showed unsurprisingly higher item decoding during encoding, this was not the case during maintenance in either impulse response (Fig. 2C bottom row).

### Stable WM coding scheme in time

The relationship between orientations and impulses/time is visualized in state-space through MDS (Fig. 4A). While the first dimension clearly differentiates between impulses, the second and third dimensions code the circular geometry of orientations in both impulses, suggesting that while the impulse responses are different between impulses, the orientation coding schemes revealed by the impulse are the same. This is corroborated by significant decoding accuracy of the impulse (*p* < 0.001, one-sided; Fig. 4B) on the one hand, but also significant cross-generalization of the orientation code between impulses (*p* < 0.001, two-sided), which was not significantly different from same-impulse orientation decoding (*p* = 0.581, two-sided; Fig. 4C).

**Figure 4.**
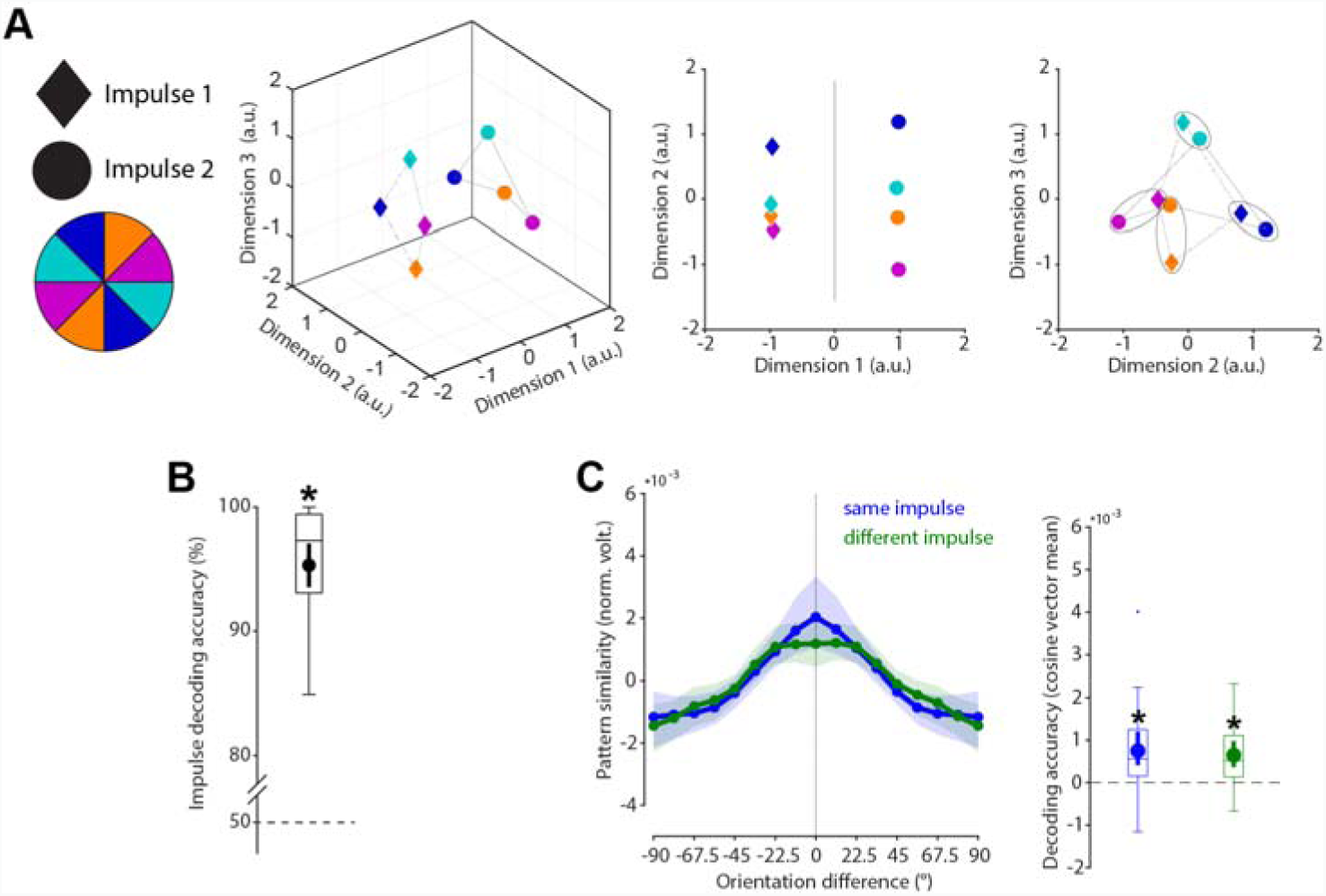
Cross-generalization of coding scheme between impulses. **(A)** Visualization of orientation and impulse code in state-space. The first dimension discriminates between impulses. The second and third dimensions code the orientation space in both impulses. **(B)** Trial-wise accuracy (%) of impulse decoding. **(C)** Orientation decoding within each impulse (blue) and orientation code cross-generalization between impulses (green). Error shadings and error bars are 95 % C.I. of the mean. Centre lines of boxplots indicate the median; box outlines show 25th and 75th percentiles, and whiskers indicate 1.5x the interquartile range. Extreme values are shown separately (dots). Asterisks indicate significant decoding accuracies or cross-generalization (p < 0.05).

It is not possible to conclude whether the difference between impulses is due to a neural network that changes during the maintenance period over time, due to different stimulation histories at the time of perturbation (i.e., the first impulse always preceded the second impulse), or due to different WM operations at each impulse event (e.g. item selection at impulse 1, response preparation at impulse 2).

To rule out that the difference in impulse response reported above is not only due to difference in stimulation history and changing WM operations, but also due to temporal coding in the WM network, we reanalysed previously published data where a single impulse stimulus was presented either 1,170 or 1,230 ms after the presentation of a single memory item (17). The findings largely replicate the results reported above: State-space visualization of impulse-onset and orientations shows the same circular geometry of the orientations at each impulse onset, while also highlighting a separation of impulse onsets in state-space (Suppl. fig. 2A). Decoding impulse-onset was significantly than from chance (*p* = 0.005, one-sided; Suppl. fig. 2B). Cross-generalization of the orientation code between impulse-onsets was significant (*p* < 0.001, two-sided), and did not significantly differ from decoding the memorized orientation within the same impulse-onset (*p* = 0.244, two-sided; Suppl. fig. 2C).

Overall, the results of the current study, as well as the reanalyses of (17) provide evidence for a low-dimensional change over time, that can be revealed by perturbing the WM network at different time-points (as predicted in (33)), while at the same time providing evidence for a temporally stable coding scheme of WM content (3,4).

### Specific WM coding scheme in space

As a counterpart to the stable coding scheme in time reported above, we explicitly tested if the coding scheme is location specific (i.e., dependent on the previous presentation location of the cued orientation). State-space visualization of cued item location and orientations shows a clear separation between locations and no overlap in orientation coding between locations (Fig. 5A). The cued location was significantly decodable from the impulse responses (*p* < 0.001, one-sided; Fig. 5B). Cross-generalization of the orientation coding scheme between cued item locations was not significant (*p* = 0.403, two-sided), and significantly lower than same side orientation decoding (*p* = 0.009, two-sided; Fig. 5C). These results reflect previous reports of spatially specific WM codes, even when location is no longer relevant (34).

**Figure 5.**
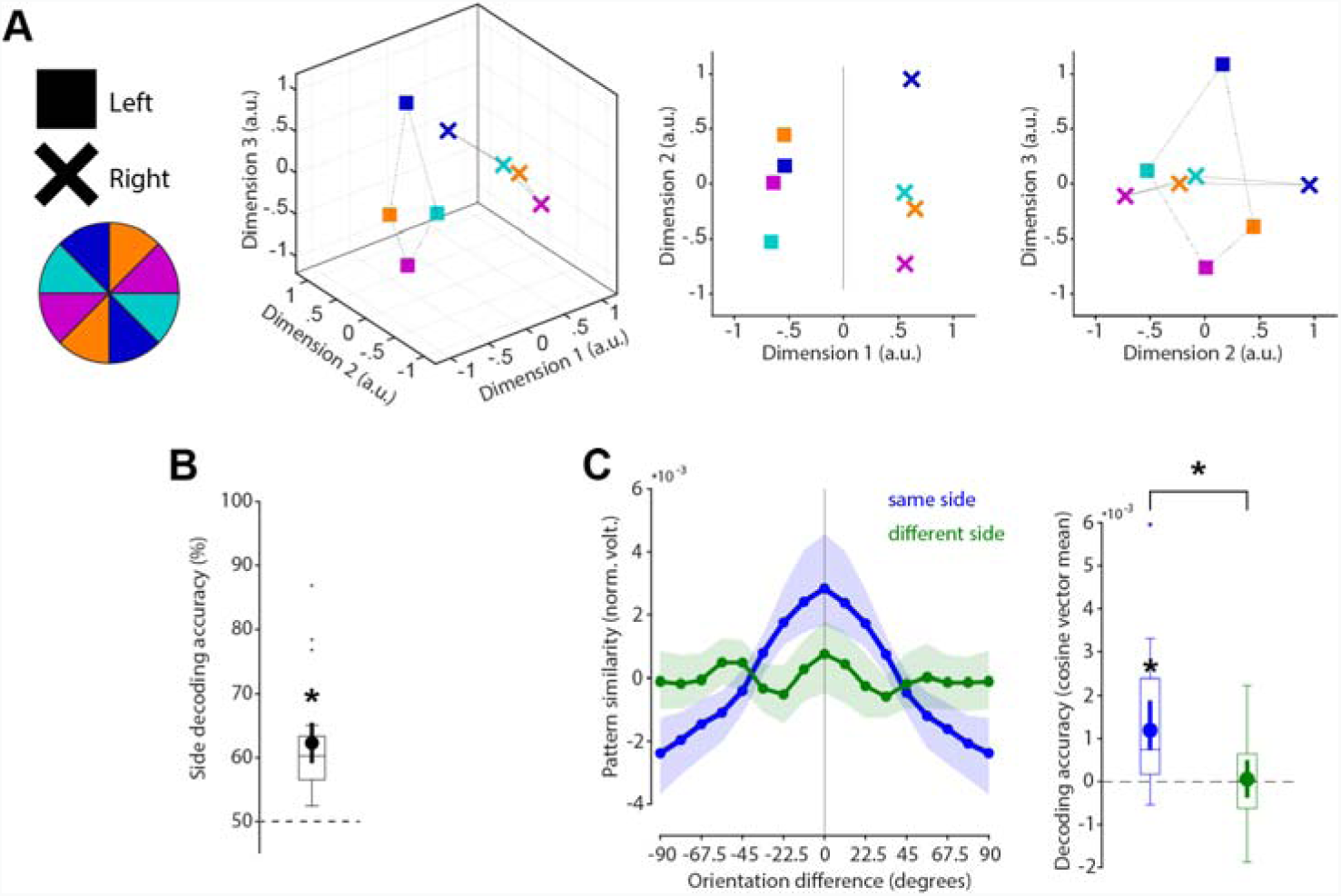
No cross-generalization of coding scheme between cued item locations during impulse responses **(A)** Visualization of orientation and item location code in state-space. The first dimension discriminates between item locations. The first and second dimensions code the orientation space, separately for WM items previously presented on the left or right side. **(B)** Trial-wise accuracy (%) of item location decoding. **(C)** Orientation decoding within each item location (blue) and orientation code cross-generalizing between different item locations (green). Error shadings and error bars are 95 % C.I. of the mean. Centre lines of boxplots indicate the median; box outlines show 25th and 75th percentiles, and whiskers indicate 1.5x the interquartile range. Extreme values are shown separately (dots). Asterisks indicate significant decoding accuracies and differences (p < 0.05).

### Drifting WM code

The first approach to test for a possible shift of the neural representation towards the response averaged the trial-wise orientation tuning curves obtained from the cross-validated orientation reconstruction on all trials (see Methods and Fig. 6A).

**Figure 6.**
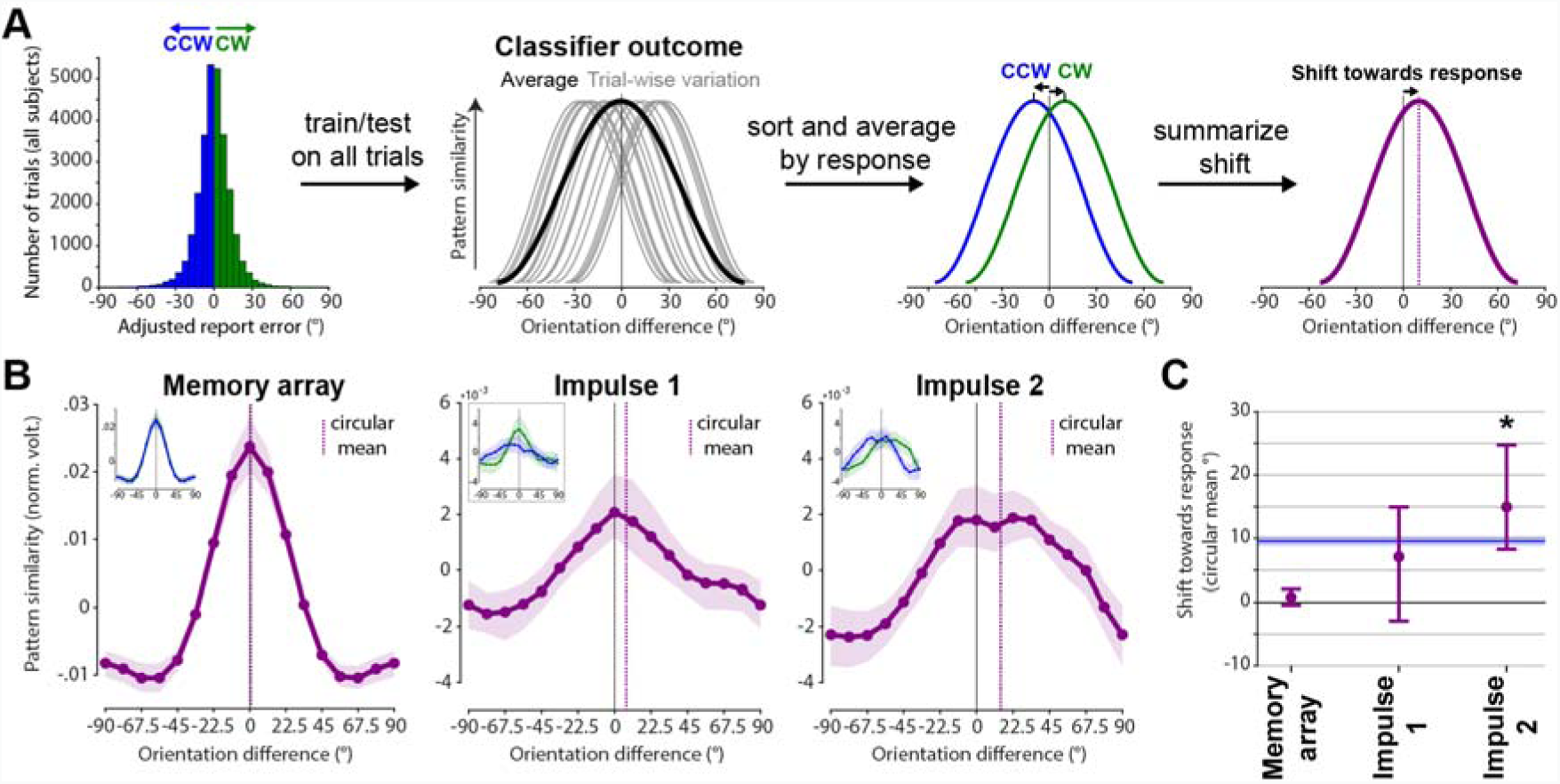
Response-dependent averaging of trial-wise tuning curves demonstrates drift. Schematic and results. **(A)** Testing for shift towards response by averaging trial-wise tuning curves by CCW/CW responses. **(B)** Results of schematised approach in A. Orientation tuning curves averaged by response such that a right-ward shift reflects a shift towards the response (purple) at each event. Purple vertical lines show circular means of the tuning curves. Insets show orientation tuning curves for CCW (blue) and CW (green) responses separately. Error shadings are 95 % C. I. of the mean. **(C)** Group-level shifts towards the response (circular mean) of each response-dependent tuning curve. Error-bars are 95 % C. I. of the mean. The blue line and shading indicates the mean and 95 % C.I. of the absolute, bias-adjusted behavioural response deviation.

No significant shift towards the response was evident during encoding/memory array presentation (*p* = 0.117, one-sided; Fig. 6B & C, left). No evidence for such a shift was found at impulse 1/early maintenance either (*p* = 0.07, one-sided; Fig. 6B & C, middle). However, the orientation tuning curve was significantly shifted towards the response at impulse 2/late maintenance (*p* < 0.001, one-sided; Fig. 6B & C, right).

The second approach to test for a possible shift of the neural representation towards the response may be more sensitive since it trains the orientation classifier only on CCW trials, and tests it on CW trials, and vice versa (see Methods and Fig. 7A), thus increasing any response related shift by a factor of two.

**Figure 7.**
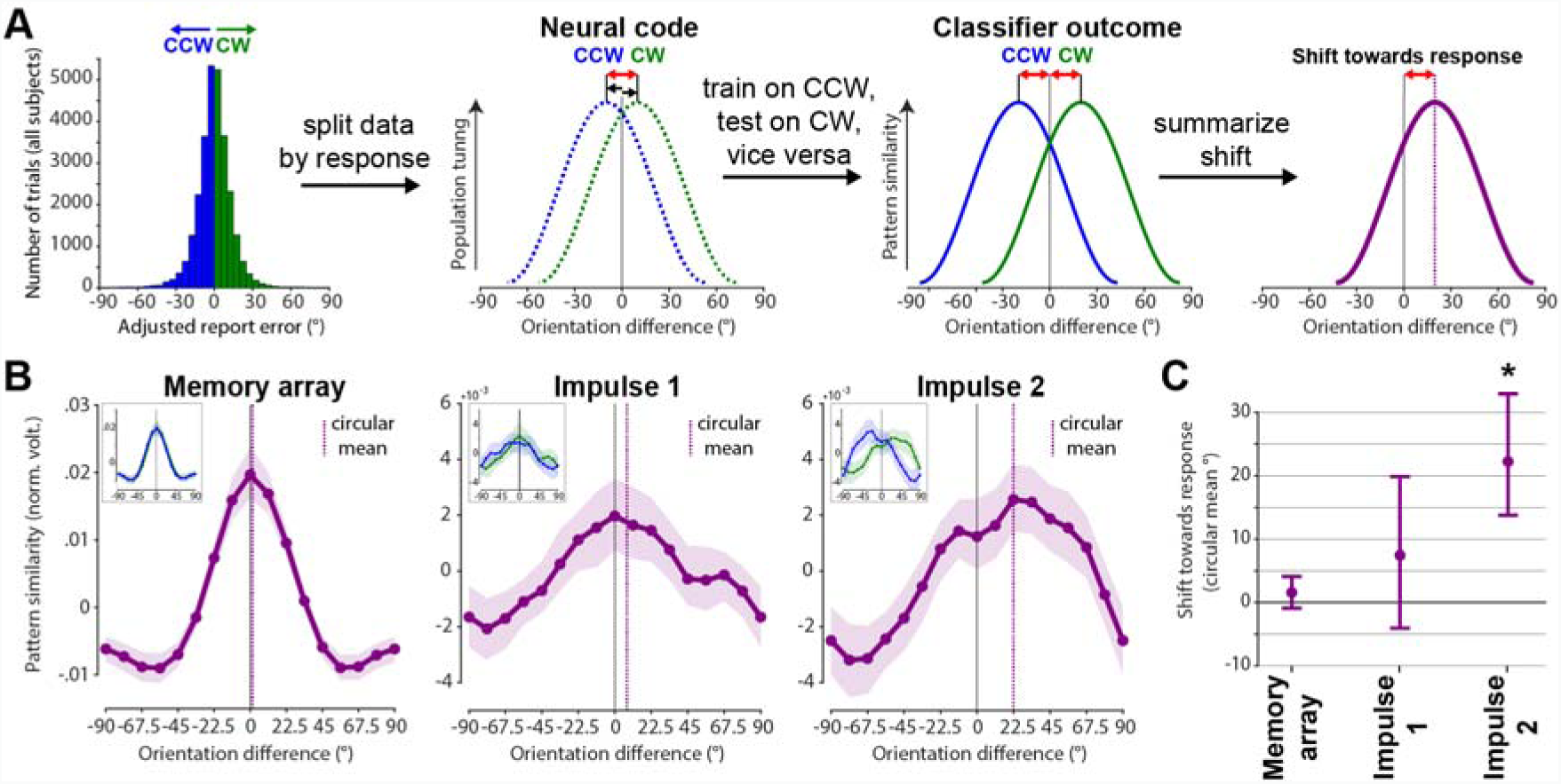
Response-dependent training and testing demonstrates drift. Schematic and results. **(A)** Testing for shift towards response by first splitting the neuroimaging data into CW and CCW data sets, and training on CW trials and testing on CCW trials, and vice versa. Given an actual shift, the shift of the resulting orientation reconstruction will be doubled, since training and testing data are shifted in opposite directions. **(B)** Results of schematised approach in A. Average orientation tuning curves such that a rightward shift reflects a shift towards the response (purple) at each event. Purple vertical lines show circular means of the tuning curves. Insets show orientation tuning curves for CCW (blue) and CW (green) responses separately. Error shadings are 95 % C. I. of the mean. **(C)** Group-level shifts towards the response (circular mean) of each response-dependent tuning curve. Error-bars are 95 % C. I. of the mean.

This approach yielded similar results as the previous approach, though the shift magnitudes are indeed larger. Neither the memory array presentation/encoding, nor impulse 1/early maintenance showed a significant shift towards the response (*p* = 0.124, *p* = 0.104, respectively, one-sided; Fig. 7, left & middle), while impulse 2/late maintenance did (*p* < 0.001, one-sided; Fig. 7, right).

Note the reported results of shifts during impulse presentations were obtained by training the classifier on both impulses, but testing it on each impulse separately. This was done to improve power (as explained in Methods). This improved orientation reconstruction particularly for the latter shift-analysis where the classifier is trained on only half the trials (CW trials only or CCW trials only). However, the same analyses based on training (and testing) within each impulse epoch separately yielded qualitatively similar results (no significant shifts at impulse 1 in either approach, significant shifts at impulse 2 in both approaches; Suppl. fig. 3).

## Discussion

In the present study, we investigated the neural dynamics of WM by probing the coding scheme over time, as well as drift in the actual memories. The neural response to impulse stimuli in this non-spatial WM paradigm enabled us to show that the coding scheme of parametric visual feature (i.e., orientation) in WM remained stable during maintenance, reflected in the significant cross-generalization of the orientation decoding between early and late impulses (Fig. 4). However, memories drift within this stable coding scheme, leading to a bias in memories (Figs. 6 and 7).

This is consistent with previous reports of a stable subspace for WM maintenance (4,5), and provides evidence for a time-invariant coding scheme for orientations maintained in WM. However, more dynamic schemes have also been reported. For example, during the early transition between encoding and maintenance (35,36). At the extreme end, some have proposed that WM could be maintained in a dynamical system, where activity evolves along a complex trajectory in neural state space (e.g. 37). Although this complicates readout (discrimination boundaries at one time point do not generalise to other time-points), such coding schemes evolve naturally from recurrent neural networks. Moreover, such dynamics also provide additional information, such as elapsed time. In the current study, we find evidence for a hybrid model (3,4): stable decoding of WM, despite dynamic activity over time.

Specifically, while there was no cost of cross-generalizing the orientation code between impulses, there was nevertheless a clear difference in the neural pattern between them, suggesting that a separate dynamic neural pattern codes the passage of time. A reanalysis of the data of a previously published study (17) confirmed these results, suggesting that the low-dimensional dynamics code for time per se (rather than impulse number). The significant decodability of impulse onset shows that the WM network changes during the maintenance even within 60 ms, resulting in distinct neural impulse responses at different time-points providing evidence for a neural time-code. Importantly, the low-dimensional representation of elapsed time is orthogonal to the mnemonic subspace, allowing WM representations to be stable. This hybrid of stable and dynamic representations may emerge from interactions between dynamic recurrent neural networks and stable sensory representations (3).

Our index of WM-related neural activity was based on an impulse response approach that we previous developed to measure WM-related changes in the functional state of the system, including ‘activity-silent’ WM states (17,18,38,39). For example, activity states during encoding could result in a neural trace in the WM network through short-term synaptic plasticity resulting in a stable code for maintenance, whereas the time-dimension could be represented in its gradual fading (33,40–42). The stable WM-content coding scheme could also be achieved by low-level activity states that self-sustain a stable code through recurrent connections, a key feature of attractor models of WM (1,43), while dynamic activity patterns are coded in an orthogonal subspace that represents time. While we did not explicitly consider tonic delay activity, it is nonetheless possible that the impulse responses also reflect non-linear interactions with low-level, persistent activity states that are otherwise difficult to measure with EEG. Therefore, we cannot rule out a contribution of persistent activity in the stable coding scheme observed here.

We also found evidence that the orientation code itself drifts along the orientation dimension, predicting recall errors. While there was no bias in the neural orientation representation at either encoding or early maintenance, the second impulse towards the end of the maintenance period revealed a code that was shifted towards the direction of response error. This pattern of results is consistent with the drift account of WM, where neural noise leads to an accumulation of error during maintenance, resulting in a still sharp, but shifted (i.e. slightly wrong) neural representation of the maintained information (1,14). While previous neurophysiological recordings from monkey PFC found evidence for drift for spatial information (15), we could demonstrate a shifting representation that more faithfully represents non-spatial WM content that is unrelated to sustained spatial attention or motor preparation, by using lateralized orientations in the present study.

Bump attractors have been proposed as an ideal neural mechanism for the maintenance of continuous representations (i.e. space, orientation, colour), where a specific feature is represented by the persistent activity “bump” of the neural population at the feature’s location along the network’s continuous feature space. Neural noise randomly shifts this bump along the feature dimension, while inhibitory and excitatory connections maintain the same overall level of activity and shape of the neural network (44,45). Random walk along the feature dimension is thus a fundamental property of bump attractors, and has been found to explain neurophysiological findings (15). Typically, this is considered within the framework of persistent working memory, however transient bursts of activity could also follow similar attractor dynamics (46,47). For example, the temporary connectivity changes of the memorized WM item may indeed slowly dissolve and become coarser, periodic activity bursts may keep this to a minimum, by periodically reinstating a sharp representation. However, since this refreshing depends on the read-out of a coarse representation, the resulting representation may be slightly wrong and thus shifted. This interplay between decaying silent WM-states that are readout and refreshed by active WM-states also predicts a drifting WM code, without depending on an unbroken chain of persistent neural activity.

Moreover, the representational drift does not necessarily have to be random. Modelling of report errors in a free recall colour WM task suggests that an increase of report errors over time may be due to separable attractor dynamics, with a systematic drift towards stable colour representations, resulting in a clustering of reports around specific colour values, in addition to random drift elicited by neural noise (13). The report bias of oblique orientations seen in the present study could be explained by a similar drift towards specific orientations, which would predict an increase of report bias for longer retention periods. However, clear behavioural evidence for such an increase in systemic report errors of orientations is lacking (10). In the present study we isolated random from systematic errors, both as a methodological necessity, but also to be able to conclude that any observed shift is due to random errors. Thus, while a systematic drift towards specific orientations might be possible, the shift in representation reported here is unrelated to it.

Our results suggest that maintenance in WM is dynamic, although the fundamental coding scheme remains stable over time. Low-dimensional dynamics could provide a valuable readout of elapsed time, whilst allowing for a time-general readout scheme for the WM content. We also show that drift within this stable coding scheme could explain loss of memory precision over time.

## Acknowledgments

This research was in part funded by a James S. McDonnell Foundation Scholar Award (220020405) and an ESRC grant (ES/S015477/1) to MGS, and by the NIHR Oxford Health Biomedical Research Centre. The Wellcome Centre for Integrative Neuroimaging is supported by core funding from the Wellcome Trust (203139/Z/16/Z). The views expressed are those of the authors and not necessarily those of the National Health Service, the National Institute for Health Research or the Department of Health. EGA is in part funded by an Open Research Area grant (464.18.114). We would like to thank N.E. Myers and D. Trübutschek for helpful comments.

## Appendix

**Supplemental figure 1.**
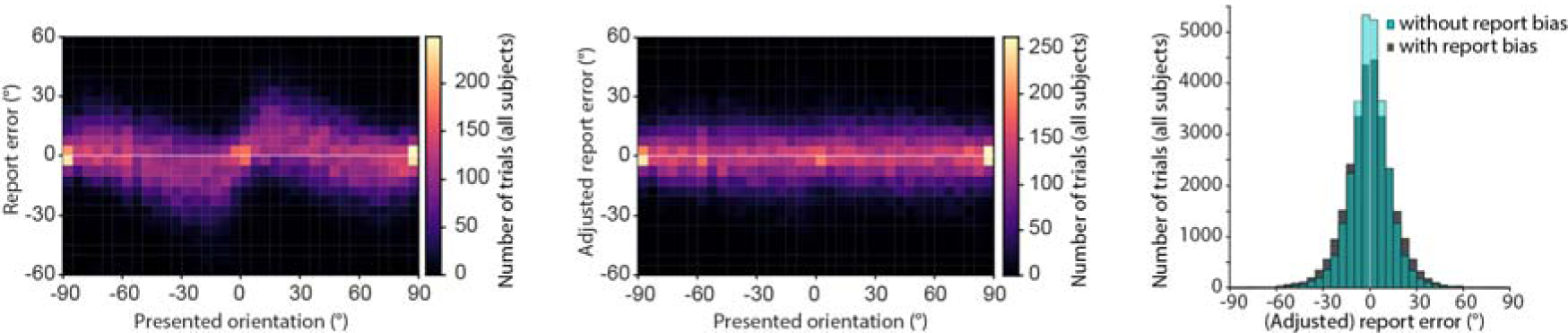
Report-bias of orientations. Participants showed a bias, exaggerating the tilt of oblique orientations, manifesting itself as a repulsion form the cardinal axes (0 and 90 degrees; *left*), similar to previous reports (32). To ensure an unbiased estimate of a possible shift in our analysis, and to isolate random from systematic errors, the report bias was removed by subtracting the median error within 11.25 degree orientation bins (*middle*). By removing orientation-specific error, the resulting error distribution is narrower (*right*). Clockwise and counter-clockwise reports were defined as positive and negative reports relative to this “adjusted”, unbiased, report error.

**Supplemental figure 2.**
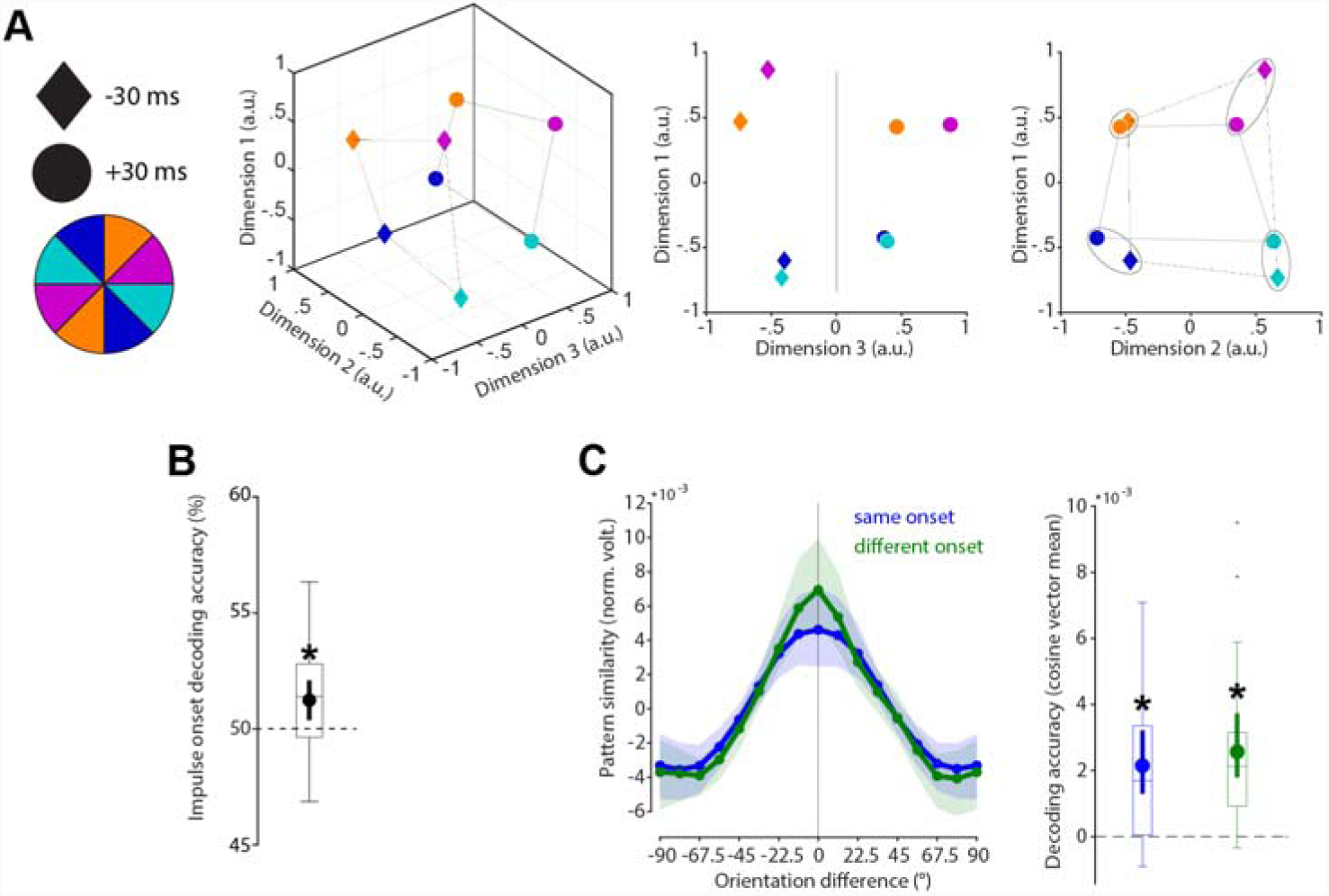
Cross-generalization of coding scheme between impulse onsets in reanalyses of (17). **(A)** Visualization of orientation and impulse-onset code in state-space. The third dimension discriminates between impulse-onsets. The first and second dimensions code the orientation space in both impulses. **(B)** Trial-wise accuracy (%) of impulse-onset decoding. **(C)** Orientation decoding within each impulse-onset (blue) and orientation code cross-generalizing between impulse-onsets (green). Error shadings and error bars are 95 % C.I. of the mean. Centre lines of boxplots indicate the median; box outlines show 25th and 75th percentiles, and whiskers indicate 1.5x the interquartile range. Extreme values are shown separately (dots). Asterisks indicate significant decoding accuracies or cross-generalization (p < 0.05).

**Supplemental figure 3.**
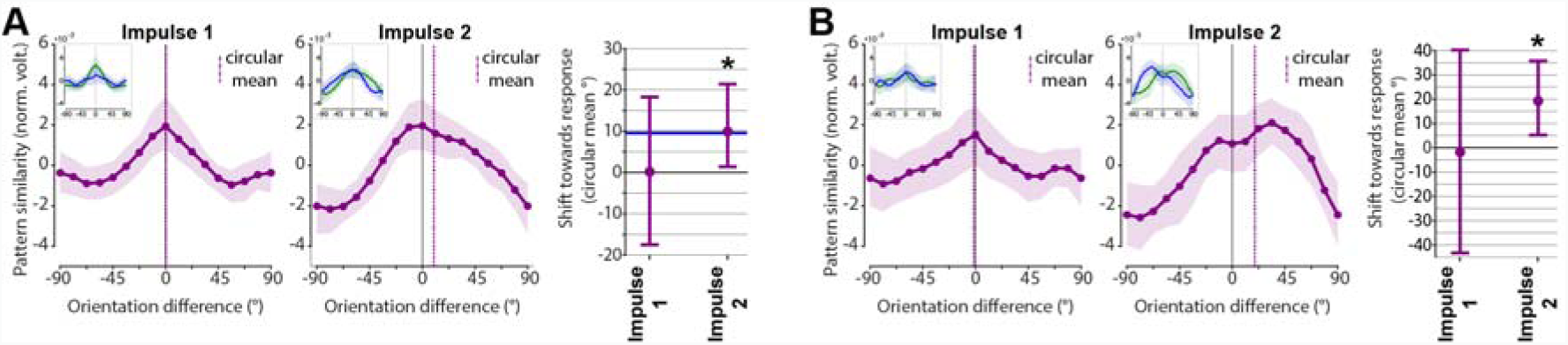
Within impulse training and testing to estimate drift. **(A)** Response-dependent averaging of trial-wise tuning curves (Fig. 6A). Shift towards response: Impulse 1: *p* = 0.4918; Impulse 2: *p* = 0.022, one-sided. **(B)** Response-dependent training and testing (Fig. 7A). Shift towards response: Impulse 1: *p* = 0.545; Impulse 2: *p* = 0.009, one-sided. Same convention as Fig. 6B-C and Fig. 7B-C.

## References

1. Compte A, Brunel N, Goldman-Rakic PS, Wang X-J. Synaptic Mechanisms and Network Dynamics Underlying Spatial Working Memory in a Cortical Network Model. Cereb Cortex. 2000 Sep 1;10(9):910–23.

2. Wang X-J. Synaptic reverberation underlying mnemonic persistent activity. Trends in Neurosciences. 2001 Aug 1;24(8):455–63.

3. Bouchacourt F, Buschman TJ. A Flexible Model of Working Memory. Neuron. 2019 Jul 3;103(1):147–160.e8.

4. Murray JD, Bernacchia A, Roy NA, Constantinidis C, Romo R, Wang X-J. Stable population coding for working memory coexists with heterogeneous neural dynamics in prefrontal cortex. PNAS. 2017 Jan 10;114(2):394–9.

5. Spaak E, Watanabe K, Funahashi S, Stokes MG. Stable and Dynamic Coding for Working Memory in Primate Prefrontal Cortex. J Neurosci. 2017 Jul 5;37(27):6503–16.

6. Cueva CJ, Marcos E, Saez A, Genovesio A, Jazayeri M, Romo R, et al. Low dimensional dynamics for working memory and time encoding. bioRxiv. 2019 Jan 31;504936.

7. Barak O, Sussillo D, Romo R, Tsodyks M, Abbott LF. From fixed points to chaos: Three models of delayed discrimination. Progress in Neurobiology. 2013 Apr 1;103:214–22.

8. Romo R, Brody CD, Hernández A, Lemus L. Neuronal correlates of parametric working memory in the prefrontal cortex. Nature. 1999 Jun;399(6735):470.

9. Druckmann S, Chklovskii DB. Neuronal Circuits Underlying Persistent Representations Despite Time Varying Activity. Current Biology. 2012 Nov 20;22(22):2095–103.

10. Rademaker RL, Park YE, Sack AT, Tong F. Evidence of gradual loss of precision for simple features and complex objects in visual working memory. Journal of Experimental Psychology: Human Perception and Performance. 2018;44(6):925–40.

11. Barrouillet P, Camos V. Developmental Increase in Working Memory Span: Resource Sharing or Temporal Decay? Journal of Memory and Language. 2001 Jul 1;45(1):1–20.

12. Kinchla RA, Smyzer F. A diffusion model of perceptual memory. Perception & Psychophysics. 1967 Jun 1;2(6):219–29.

13. Panichello MF, DePasquale B, Pillow JW, Buschman TJ. Error-correcting dynamics in visual working memory. Nature Communications. in press;

14. Schneegans S, Bays PM. Drift in Neural Population Activity Causes Working Memory to Deteriorate Over Time. J Neurosci. 2018 May 23;38(21):4859–69.

15. Wimmer K, Nykamp DQ, Constantinidis C, Compte A. Bump attractor dynamics in prefrontal cortex explains behavioral precision in spatial working memory. Nat Neurosci. 2014 Mar;17(3):431–9.

16. Lim PC, Ward EJ, Vickery TJ, Johnson MR. Not-so-working Memory: Drift in Functional Magnetic Resonance Imaging Pattern Representations during Maintenance Predicts Errors in a Visual Working Memory Task. Journal of Cognitive Neuroscience. 2019 May 21;1–15.

17. Wolff MJ, Ding J, Myers NE, Stokes MG. Revealing hidden states in visual working memory using electroencephalography. Front Syst Neurosci. 2015

18. Wolff MJ, Jochim J, Akyürek EG, Stokes MG. Dynamic hidden states underlying working-memory-guided behavior. Nature Neuroscience. 2017 Jun;20(6):864–71.

19. Kleiner M. Visual stimulus timing precision in Psychtoolbox-3: Tests, pitfalls solutions [Internet]. 2010 [cited 2016 Jul 12]. Available from: http://www.neuroschool-tuebingen-nena.de/fileadmin/user-upload/Dokumente/neuroscience/AbstractbookNeNa2010u.pdf

20. Delorme A, Makeig S. EEGLAB: an open source toolbox for analysis of single-trial EEG dynamics including independent component analysis. Journal of Neuroscience Methods. 2004 Mar;134(1):9–21.

21. Hyvarinen A. Fast and robust fixed-point algorithms for independent component analysis. IEEE Transactions on Neural Networks. 1999 May;10(3):626–34.

22. Fritsche M, Mostert P, de Lange FP. Opposite Effects of Recent History on Perception and Decision. Current Biology. 2017 Feb 20;27(4):590–5.

23. Grootswagers T, Wardle SG, Carlson TA. Decoding Dynamic Brain Patterns from Evoked Responses: A Tutorial on Multivariate Pattern Analysis Applied to Time Series Neuroimaging Data. J Cogn Neurosci. 2017 Apr;29(4):677–97.

24. Nemrodov D, Niemeier M, Patel A, Nestor A. The Neural Dynamics of Facial Identity Processing: Insights from EEG-Based Pattern Analysis and Image Reconstruction. eNeuro. 2018 Jan 1;5(1):ENEURO.0358-17.2018.

25. Wolff MJ, Kandemir G, Stokes MG, Akyurek EG. Impulse responses reveal unimodal and bimodal access to visual and auditory working memory. bioRxiv. 2019 Apr 30;623835.

26. Ledoit O, Wolf M. Honey, I shrunk the sample covariance matrix. The Journal of Portfolio Management. 2004;30(4):110–9.

27. Myers NE, Rohenkohl G, Wyart V, Woolrich MW, Nobre AC, Stokes MG. Testing sensory evidence against mnemonic templates. eLife. 2015 Dec 14;4:e09000.

28. Serences JT, Saproo S. Computational advances towards linking BOLD and behavior. Neuropsychologia. 2012 Mar 1;50(4):435–46.

29. Brouwer GJ, Heeger DJ. Decoding and reconstructing color from responses in human visual cortex. J Neurosci. 2009 Nov 4;29(44):13992–4003.

30. Ede F van, Chekroud SR, Stokes MG, Nobre AC. Concurrent visual and motor selection during visual working memory guided action. Nature Neuroscience. 2019 Mar;22(3):477.

31. Oostenveld R, Fries P, Maris E, Schoffelen J-M, Oostenveld R, Fries P, et al. FieldTrip: Open Source Software for Advanced Analysis of MEG, EEG, and Invasive Electrophysiological Data, FieldTrip: Open Source Software for Advanced Analysis of MEG, EEG, and Invasive Electrophysiological Data. Computational Intelligence and Neuroscience, Computational Intelligence and Neuroscience. 2010 Dec 23;2011, 2011:e156869.

32. Pratte MS, Park YE, Rademaker RL, Tong F. Accounting for stimulus-specific variation in precision reveals a discrete capacity limit in visual working memory. Journal of Experimental Psychology: Human Perception and Performance. 2017;43(1):6–17.

33. Buonomano DV, Maass W. State-dependent computations: spatiotemporal processing in cortical networks. Nature Reviews Neuroscience. 2009 Feb;10(2):113–25.

34. Pratte MS, Tong F. Spatial specificity of working memory representations in the early visual cortex. Journal of Vision. 2014 Mar 1;14(3):22–22.

35. Wasmuht DF, Spaak E, Buschman TJ, Miller EK, Stokes MG. Intrinsic neuronal dynamics predict distinct functional roles during working memory. Nature Communications. 2018 Aug 29;9(1):3499.

36. Cavanagh SE, Towers JP, Wallis JD, Hunt LT, Kennerley SW. Reconciling persistent and dynamic hypotheses of working memory coding in prefrontal cortex. Nature Communications. 2018 Aug 29;9(1):3498.

37. Maass W, Natschläger T, Markram H. Real-time computing without stable states: a new framework for neural computation based on perturbations. Neural Comput. 2002 Nov;14(11):2531–60.

38. Stokes MG. ‘Activity-silent’ working memory in prefrontal cortex: a dynamic coding framework. Trends in Cognitive Sciences. 2015 Jul;19(7):394–405.

39. Masse NY, Yang GR, Song HF, Wang X-J, Freedman DJ. Circuit mechanisms for the maintenance and manipulation of information in working memory. Nature Neuroscience. 2019 Jun 10;1.

40. Nikolić D, Häusler, Stefan, Singer, Wolf, Maass, Wolfgang. Temporal dynamics of information content carried by neurons in the primary visual cortex. Advances in Neural Information Processing Systems. 2007;19:1041–1048.

41. Nikolić D, Häusler S, Singer W, Maass W. Distributed Fading Memory for Stimulus Properties in the Primary Visual Cortex. PLOS Biol. 2009 Dec 22;7(12):e1000260.

42. Zucker RS, Regehr WG. Short-Term Synaptic Plasticity. Annu Rev Physiol. 2002 Mar 1;64(1):355–405.

43. Chaudhuri R, Fiete I. Computational principles of memory. Nature Neuroscience. 2016 Mar;19(3):394–403.

44. Amari S. Dynamics of pattern formation in lateral-inhibition type neural fields. Biol Cybern. 1977 Jun 1;27(2):77–87.

45. Brody CD, Romo R, Kepecs A. Basic mechanisms for graded persistent activity: discrete attractors, continuous attractors, and dynamic representations. Current Opinion in Neurobiology. 2003 Apr 1;13(2):204–11.

46. Lundqvist M, Herman P, Lansner A. Theta and Gamma Power Increases and Alpha/Beta Power Decreases with Memory Load in an Attractor Network Model. Journal of Cognitive Neuroscience. 2011 Mar 31;23(10):3008–20.

47. Mongillo G, Barak O, Tsodyks M. Synaptic Theory of Working Memory. Science. 2008 Mar 14;319(5869):1543–6.

